# Dopaminergic neurons are vulnerable to dysregulation of YEATS2-dependent calcium homeostasis

**DOI:** 10.1101/2025.09.06.674642

**Authors:** Luca Lo Piccolo, Ranchana Yeewa, Pitiporn Noisagul, Arnaud Monteil, Vorasuk Shotelersuk, Salinee Jantrapirom

## Abstract

YEATS2 is a ubiquitously expressed chromatin-associated factor that we recently identified as a novel regulator of dopaminergic (DAergic) synaptic integrity, though its mechanism of action remained unclear. Using *Drosophila*, we show that neuronal depletion of YEATS2 reshapes the brain transcriptome, marked by downregulation of metabolic genes and upregulation of G protein–coupled receptors (GPCRs). These changes coincide with elevated intracellular calcium, neurobehavioral deficits, and selective DA neuron loss. Importantly, genetic or pharmacological inhibition of the store-operated calcium entry channel Orai restored calcium homeostasis and rescued DA neuron survival. Our findings define a YEATS2-dependent epigenetic–calcium axis that governs DA neuron vulnerability.

## Introduction

YEATS2 is a chromatin-associated protein that selectively recognizes histone crotonylation, recruiting chromatin remodelling complexes and transcriptional co-factors to regulate gene expression programs essential for cellular differentiation and homeostasis (Orpinell et al. 2010; Suganuma et al. 2010; Zhao et al. 2016). While extensively studied in cancer, especially solid tumours, YEATS2’s roles in non-malignant tissues remain largely unexplored (Roy et al. 2025; Zhai et al. 2025). Our recent work in *Drosophila* showed that neuron-specific knockdown of *dYEATS2* disrupts dopaminergic (DAergic) synaptic integrity and induces seizure-like behaviours, highlighting YEATS2 as a key regulator of neuronal stability (Lo Piccolo et al. 2024).

DAergic neurons govern motor control, reward processing, and a broad spectrum of cognitive functions, integrating signals across neural circuits underlying voluntary movement, motivation, and learning (Puig and Miller 2012; Ott et al. 2023). Despite their importance, these neurons exhibit selective vulnerability in Parkinson’s disease (PD), where their progressive loss leads to debilitating symptoms (Surmeier et al. 2017). Multiple stressors, including high metabolic demand, dopamine oxidation, calcium imbalance, mitochondrial dysfunction, and α-synuclein aggregation, have been implicated in this vulnerability. Yet, how these factors coalesce to drive selective DAergic neuron degeneration remains incompletely understood, impeding therapeutic advances.

Epigenetic regulation has emerged as a critical modulator of neuronal identity, plasticity, and survival. Perturbations in epigenetic mechanisms profoundly disrupt DAergic neuron function by triggering maladaptive gene expression and increasing susceptibility to stress (Murphy and Heller 2022). However, the epigenetic factors and downstream pathways that mediate DAergic neuron resilience and vulnerability remain largely undefined.

Investigating how YEATS2, as an epigenetic regulator, maintains DAergic neuron integrity could reveal key molecular mechanisms underlying their selective vulnerability and the networks that safeguard neuronal health. In this study, we address this question by characterizing the transcriptional consequences of neuronal YEATS2 depletion and assessing its impact on DAergic neuron survival.

## Results and Discussion

### *dYEATS2* knockdown induces transcriptional shifts in neuronal signalling and metabolism

To investigate the molecular consequences of neuronal *YEATS2* depletion, we performed RNA-seq on *Drosophila* heads with pan-neuronal *dYEATS2* knockdown (*dYEATS2-IR*) and compared these to *non-targeting control* (*NTC IR-1*), as well as additional in-house controls, including knockdown of the YEATS paralog *dENL/AF9* and a second *non-targeting control* (*NTC IR-2*) (Figure 1A; Table S1). Raw and DESeq2-normalized RNA-seq data have been deposited in the Gene Expression Omnibus (GSE307078). We performed gene set enrichment analysis (GSEA) to identify biological pathways significantly affected by YEATS2 depletion (Figure 1B, C). To prioritize the most robust and biologically relevant pathways, we applied stringent cutoffs, normalized enrichment score (NES) > 2 and false discovery rate (FDR q) < 0.05 (Figure 1C-E). This approach enhanced specificity and enabled us to dissect key regulatory networks perturbed by YEATS2 deficiency.

**Figure 1.**
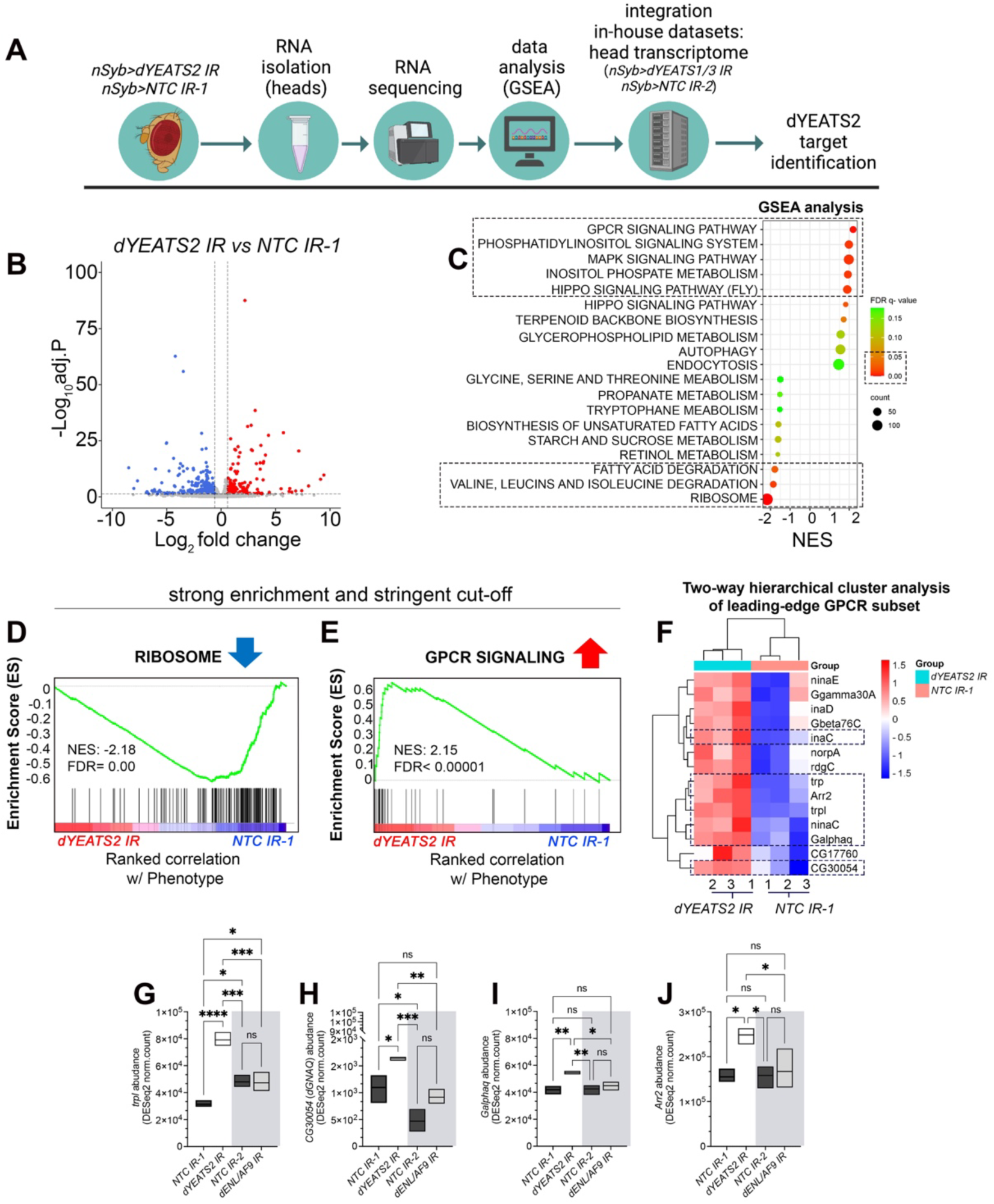
YEATS2 knockdown alters brain transcriptome with enrichment for metabolic and GPCR signalling genes. (A) Schematic of the RNA-seq workflow. Brains with pan-neuronal *dYEATS2* knockdown (*dYEATS2-IR*) were compared to a non-targeting control (*NTC IR-1*), using RNA extracted from 50 brains per replicate across three biological replicates (150 brains per group). To assess target specificity, additional comparisons were made with in-house RNA-seq datasets from *YEATS* paralog knockdowns (*dENL/AF*) and a second independent control (*NTC IR-2*). (B) Volcano plot showing differentially expressed genes (DEGs) in *dYEATS2-IR* brains, with a total of 1,160 DEGs identified (284 upregulated and 876 downregulated). Up-regulated genes (log₂FC > 0.585, FDR < 0.05) are in red; down-regulated genes (log₂FC < –0.585, FDR < 0.05) in blue; non-significant genes in gray. (C) Gene set enrichment analysis (GSEA) of DEGs. Selected gene sets with FDR q-values < 0.15 are shown. Dotted boxes highlight gene sets with FDR q < 0.05 considered for further analysis. (D, E) GSEA with stringent cutoff (normalized enrichment score [NES] > 2) identified strong enrichment for ribosome-related genes (down-regulated) and G protein–coupled receptor (GPCR) signalling genes (up-regulated). (F) Two-way hierarchical clustering of leading-edge genes in the GPCR signalling gene set. Dotted boxes highlight genes with consistent expression patterns across replicates, including *trpL*, *CG30054*, *GalphaQ*, and *Arr2*. (G–J) Normalized expression counts for selected GPCR genes in control (*dYEATS2-IR*), and additional internal controls (*NTC IR-2* and *dENL/AF9-IR*). Data show increased expression specific to *dYEATS2* knockdown.

Consistent with prior studies in non-small cell lung cancer models (Mi et al. 2017), we observed significant downregulation of genes encoding ribosomal proteins and translation machinery following *YEATS2* knockdown (Figure 1D; Figure S1). This conserved transcriptional signature across species and cell types supports an essential housekeeping role for YEATS2 in sustaining fundamental cellular processes. Given YEATS2’s established function as a component of the ATAC chromatin remodelling complex, which promotes gene activation at H3K27ac-enriched promoters, this downregulation likely reflects a direct consequence of disrupted chromatin-mediated gene activation.

Although our stringent cutoffs limited detailed analysis of metabolic genes downregulated by dYEATS2 depletion, it is noteworthy that YEATS family proteins have been implicated in regulating cellular metabolism across diverse organisms (Zhang et al. 2021; Yeewa et al. 2024; Pant et al. 2025). This evolutionary conservation, from yeast to humans, underscores their fundamental role in translating metabolic states into transcriptional and epigenetic outputs.

Alongside ribosomal gene downregulation, we detected prominent upregulation of G protein– coupled receptor (GPCR) signalling genes and phosphatidylinositol pathway components. Two-way hierarchical clustering of leading-edge genes revealed distinct grouping patterns among replicates and highlighted major gene expression changes associated with *YEATS2* knockdown (Figure 1F-J). We focused on *trpL*, *CG30054*, *GalphaQ*, and *Arr*2 for further analysis due to their consistent upregulation and known roles in neuronal GPCR signalling and calcium regulation. Specifically, *trpL* encodes a transient receptor potential (TRP)-like channel involved in calcium influx; *CG30054* is predicted to enable G protein-coupled receptor binding activity and to be involved in signal transduction pathways; *GalphaQ* is the guanine nucleotide–binding protein subunit alpha and functions as a G protein alpha subunit mediating GPCR signalling; and *arrestin 2* (*Arr2*) regulates GPCR desensitization and internalization (Hardie et al. 2001; Pfeil et al. 2020). While other genes such as *inaC* and *nina*C were also upregulated, their functions are more specialized to phototransduction and thus less directly relevant to dopaminergic neuron signalling (Yau and Hardie 2009). These expression changes were subsequently validated by RT-qPCR (Figure S2). Because YEATS2 does not function as a classical transcriptional repressor, the observed GPCR upregulation likely arises indirectly through compensatory or secondary mechanisms triggered by YEATS2 depletion and associated cellular stress. Mechanistically, this induction may represent an adaptive response to maintain neuronal homeostasis when core transcriptional programs are perturbed. More broadly, these findings redefine YEATS2 as a hub within complex cellular networks rather than a simple molecular switch, illustrating how chromatin regulators integrate epigenetic, metabolic, and signalling cues to preserve neuronal integrity. Notably, both GPCR and phosphatidylinositol signalling pathways converge on intracellular processes that modulate calcium dynamics (Brands et al. 2024), suggesting that calcium homeostasis may be disrupted as a downstream consequence.

Taken together, these coordinated transcriptional shifts reflect both downregulation of core translational machinery and upregulation of neuronal signalling pathways, highlighting YEATS2’s broad impact on gene expression programs controlling metabolism and signal transduction.

### dYEATS2 deficiency disrupts calcium homeostasis in adult *Drosophila* brains

Given that intracellular calcium is a common regulatory node for both GPCR and phosphatidylinositol signalling, the transcriptional signature induced by *dYEATS2* depletion implicates calcium homeostasis as a downstream axis connecting chromatin mis-regulation to neuronal dysfunction. To directly test this hypothesis, we expressed the genetically encoded calcium indicator GCaMP6s (Zhang et al. 2023) pan-neuronally and performed *ex vivo* imaging of adult fly brains. Brains were incubated in PBS, NaCl, KCl, or glucose, and relative calcium responses (ΔF/F₀) were quantified after normalization to the PBS baseline to account for potential differences in GCaMP expression between genotypes (Figure 2A). Under these conditions, dYEATS2-depleted neurons exhibited significantly enhanced calcium elevations in response to depolarization (KCl) and metabolic stimulation (glucose) compared to controls (Figure 2). These results indicate that dYEATS2 deficiency sensitizes neurons to physiological stimuli, reflecting impaired control of intracellular calcium flux. Mechanistically, the enhanced calcium responses are consistent with transcriptional upregulation of GPCR signalling pathway, which is well-established regulators of calcium entry. By linking *dYEATS2* depletion to exaggerated calcium dynamics, these findings underscore an epigenetic-to-physiological axis whereby disruption of chromatin regulation propagates downstream to perturb neuronal homeostasis. This state of neuronal hypersensitivity likely predisposes neurons to functional stress and hyperexcitability, offering a mechanistic explanation for the seizure-like behaviours previously observed in dYEATS2-deficient flies (Lo Piccolo et al. 2024).

**Figure 2.**
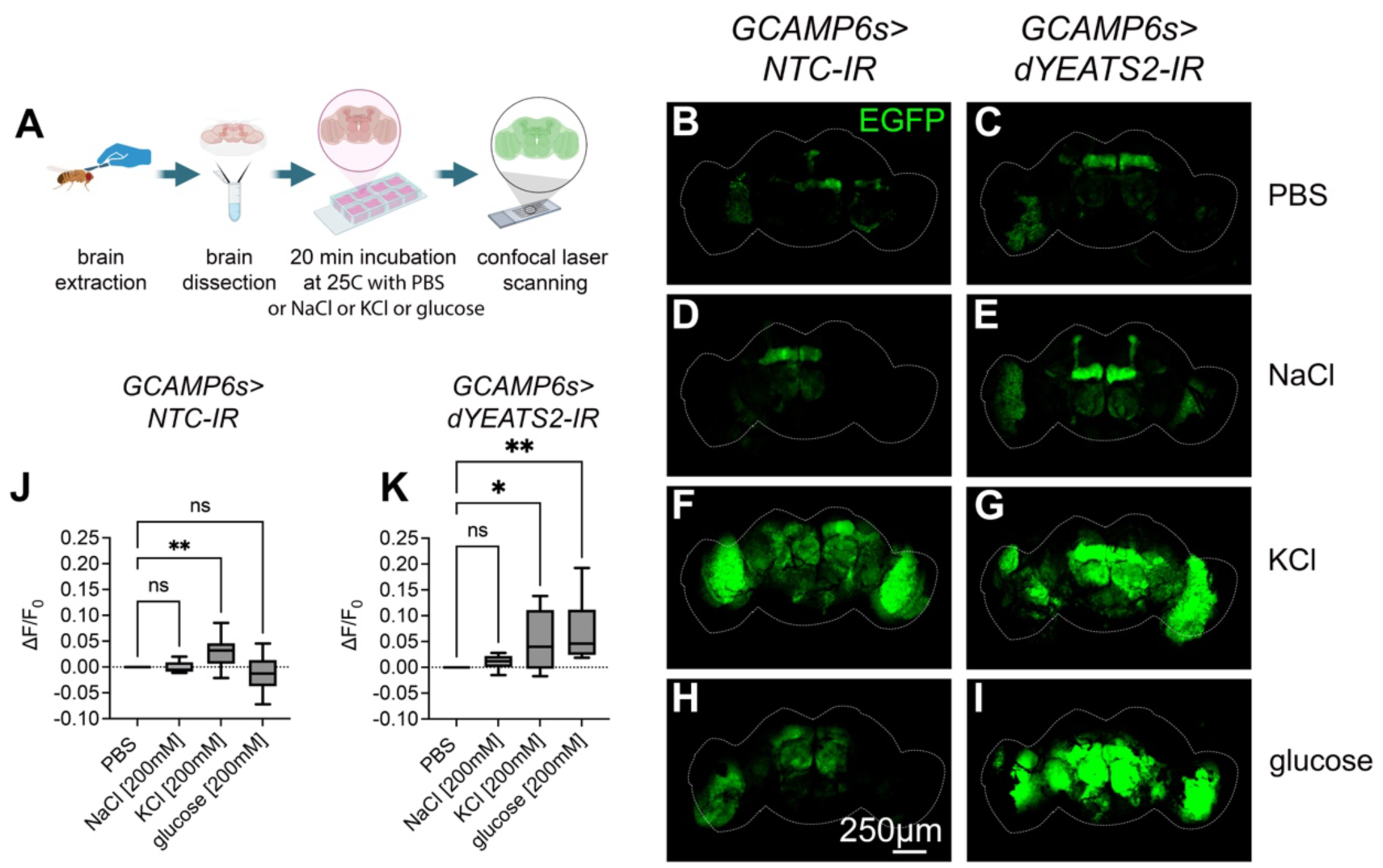
YEATS2 depletion induces calcium overload in adult fly brains, revealed by *ex vivo* GCaMP6s imaging. (A) Schematic of the experimental paradigm. Adult *Drosophila* brains expressing the genetically encoded calcium indicator GCaMP6s and *dYEATS2* RNAi under pan-neuronal control (*GCaMP6s>dYEATS2-IR*) were dissected and incubated ex vivo for 20 min at 25 °C in PBS, 200 mM NaCl, 200 mM KCl, or 200 mM glucose. Brains co-expressing GCaMP6s and a *non-targeting mCherry RNAi* (*GCaMP6s>NTC-IR*) served as controls. For each condition, 10 brains were analyzed per group. (B–I) Representative confocal images of adult brains acquired in anterior–posterior orientation, presented as maximum intensity projections from 12 optical sections (4 μm each). GCaMP6s fluorescence reflects intracellular calcium levels. (J, K) Quantification of relative calcium signals (ΔF/F₀), normalized to the PBS baseline to control for differences in GCaMP expression across genotypes, reveals significantly enhanced stimulus-evoked calcium responses in YEATS2-depleted neurons compared to controls. Increased calcium elevations upon depolarization (KCl) and metabolic stimulation (glucose) indicate aberrant calcium handling following YEATS2 loss. Data are presented as mean ± SEM. Statistical comparisons were performed using one-way ANOVA followed by Dunnett’s post hoc analysis. p < 0.05 was considered statistically significant.

To assess causality, we attempted to mitigate these calcium abnormalities. Initial targeting of Gαq, a GPCR effector upregulated upon *dYEATS2* knockdown, was lethal when co-expressed with *dYEATS2 RNAi*, highlighting the essentiality of precise GPCR signalling in neuronal viability. As an alternative, we modulated store-operated calcium entry (SOCE) via the Orai channel (Figure 3A), a key mediator of calcium influx in neurons (Feske et al. 2006). Interestingly, pan-neuronal expression of a dominant-negative Orai^E180A^ (Yeromin et al. 2006) significantly reduced seizure-like behaviours in adults (Figure 3B), while pharmacological inhibition using the pyrazole derivative BTP2 (Ohga et al. 2008) improved recovery times following electric stimulation in larvae (Figure 3C). These interventions support the hypothesis that elevated intracellular calcium functions as a critical mediator of the neurological phenotypes induced by dYEATS2 depletion.

**Figure 3.**
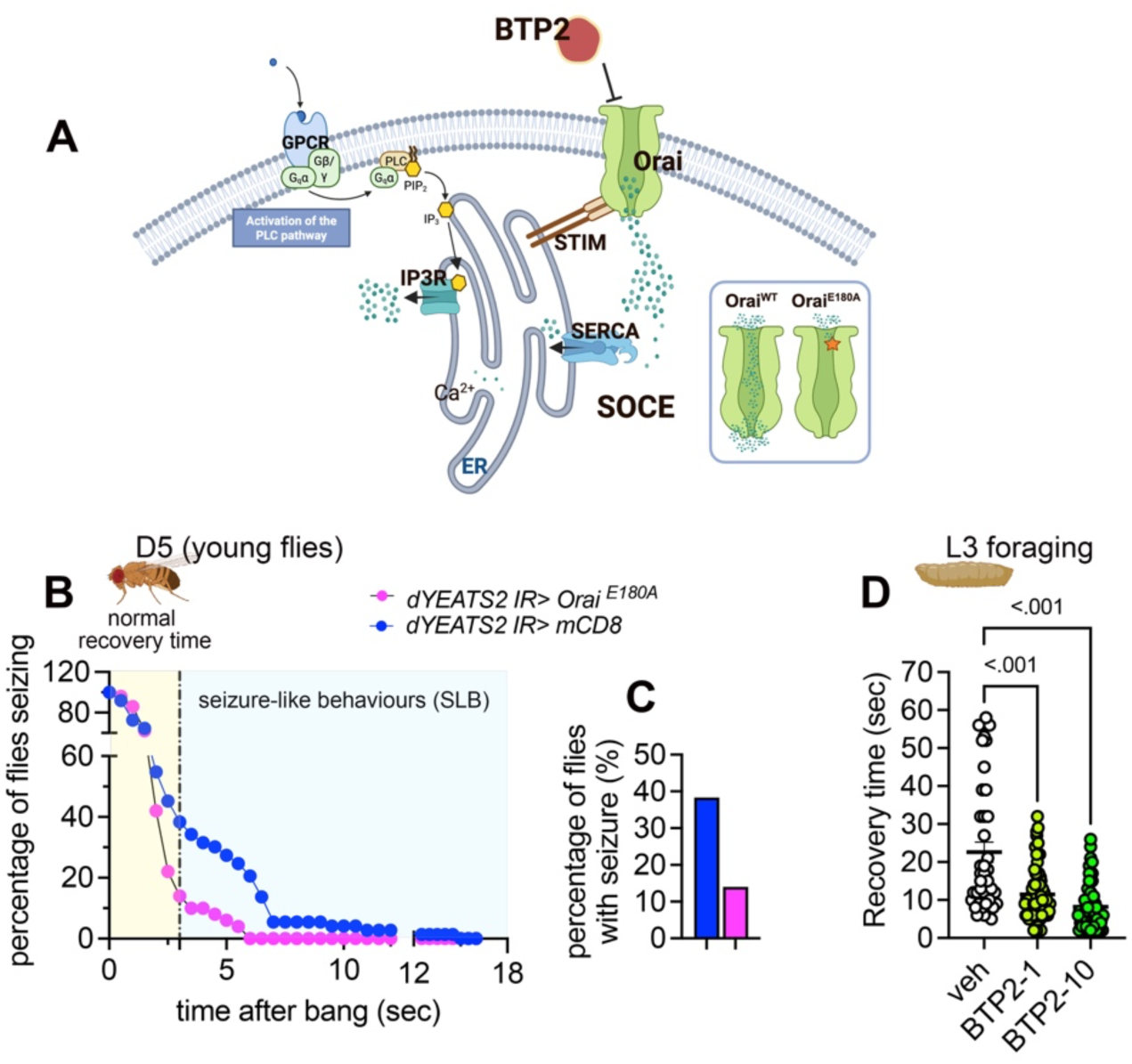
Inhibition of store-operated calcium entry (SOCE) via Orai channel ameliorates YEATS2 neurological dysfunctions. (A) Schematic model illustrating the role of Orai-mediated SOCE in GPCR-induced calcium dynamics. GPCR activation promotes intracellular calcium release from the endoplasmic reticulum (ER), followed by calcium re-entry through the Orai channel. To genetically inhibit SOCE, a dominant-negative form of Orai (*Orai180A*) was expressed pan-neuronally in the *dYEATS2* RNAi background. Pharmacological inhibition was achieved by feeding adult flies with BTP2 (YM-58483), a selective Orai channel antagonist. (B, C) Suppression of seizure-like behaviour (SLB) by Orai inhibition in adult flies. SLB was assessed at 5 days post-eclosion following mechanical stimulation (“bang” assay). Flies that recovered within 3 seconds were classified as normal; longer recovery indicated SLB. The percentage of flies exhibiting SLB was significantly reduced by genetic or pharmacological inhibition of Orai (n = 45 flies per group). (D) Pharmacological inhibition of Orai with BTP2 reduces delayed recovery after electric shock in third instar larvae. Larvae were treated with either 1 μM or 10 μM BTP2 or vehicle control, and time to recovery was measured (n = 75 larvae per group). All data are shown as mean ± SEM. Statistical significance was assessed using one-way ANOVA followed by Šidák’s post hoc testing; differences with *p* < 0.05 were considered statistically significant.

Collectively, these findings establish a causal link between epigenetic perturbation, maladaptive calcium signalling, and neuronal dysfunction, positioning YEATS2 deficiency as a novel driver of transcriptional dysregulation that promotes pathological calcium entry and neuronal hyperexcitability.

### Calcium overload resulting from *dYEATS2* knockdown drives DAergic neuron loss

Given that depletion of dYEATS2 in the nervous system reduces tyrosine hydroxylase (TH) expression and dopamine levels (Lo Piccolo et al. 2024) and having established that dYEATS2 deficiency elevates intracellular calcium and sensitizes neurons to physiological stimuli, we next asked whether this calcium perturbations compromise the integrity of DAergic neurons. To test this, we generated a stable fly line co-expressing *mCD8::EGFP* and *dYEATS2 RNAi* under pan-neuronal control, confirming proper GFP labelling and expected reductions in both *dYEATS2* and TH (Figure S3).

Analysis of DAergic neuron clusters in young adult flies revealed a pronounced loss of neurons, whereas other populations were largely preserved (Figures 4A, B and G; Figure S4). Glutamatergic neurons, which constitute most excitatory neurons in the fly brain, displayed normal cluster numbers (Figures 4C, D and H), and mushroom body architecture, representing higher-order associative circuits, was unaltered (Figures 4E, F). These observations indicate a selective vulnerability of DAergic neurons to *dYEATS2* decline, suggesting that calcium dysregulation preferentially destabilizes this population.

**Figure 4.**
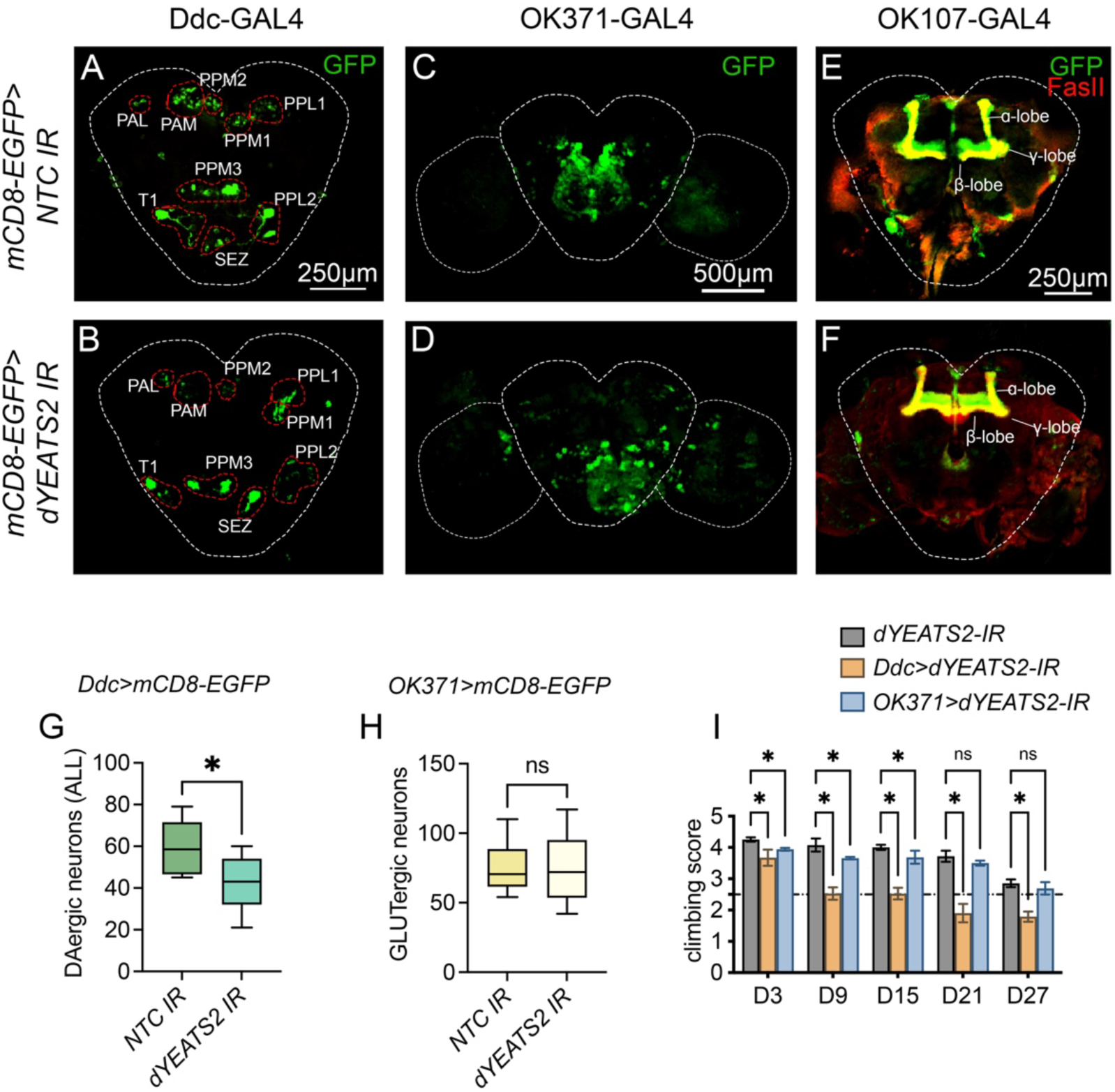
Calcium overload resulting from YEATS2 knockdown drives dopaminergic neuron loss. (A–F) Representative confocal images of distinct neuronal populations in adult *Drosophila* brains expressing membrane-bound mCD8-EGFP, driven by cell type–specific GAL4 lines, in the presence of *dYEATS2* RNAi (mCD8-EGFP > *dYEATS2*-IR) or a non-targeting RNAi control (mCD8-EGFP > NTC-IR). (A,B) Dopaminergic neurons were visualized using Ddc-GAL4. Brains are shown in posterior-to-anterior orientation; central brain regions are outlined (dotted lines), and major dopaminergic clusters, including PAL, PAM, PPL1, and PPM3, are circled in red. (C,D) Glutamatergic neurons were labelled using *OK371-GAL4*. The central brain and optic lobes are demarcated by dotted outlines. (E,F) Mushroom body neurons were visualized by FasII staining in brains expressing EGFP under *OK107-GAL4* control; α, β, and γ lobes are indicated. (G) Quantification of total dopaminergic neurons (n = 10). Additional representative brain images are provided in Figure S4. (H) Quantification of glutamatergic neurons (n = 10). (I) Age-dependent locomotor decline assessed via climbing assays in flies expressing *dYEATS2* RNAi under *Ddc-GAL4* (DAergic neurons) or *OK371-GAL4* (GLUTergic neurons) compared with genetic controls *dYEATS2 RNAi* carrying fly line lacking a GAL4 driver (non-driven control, NDC). Behavioural deficits were monitored across multiple timepoints post-eclosion (3, 9, 15, 21, and 27 days). Data represent mean ± SEM. Statistical analyses were performed using one-way ANOVA followed by Dunnett’s post hoc tests; *p* < 0.05 was considered significant.

This selective susceptibility aligns with established principles that DAergic neurons are uniquely burdened by autonomous pacemaking activity, which engages L-type Ca²⁺ channels and drives sustained calcium influx, imposing extraordinary demands on calcium buffering systems (Surmeier et al. 2011). In this context, *dYEATS2* depletion likely exacerbates chronic calcium elevation beyond neuronal buffering capacity, triggering excitotoxic stress, metabolic imbalance, and ultimately degeneration. This selective vulnerability may also have network-level consequences, given the established role of dopamine in seizure modulation: D1 receptor signalling promotes excitability, whereas D2 receptor pathways are protective (Bozzi and Borrelli 2013). Accordingly, loss of DA neurons and diminished D2 signalling in dYEATS2-depleted flies could plausibly shift the balance toward hyperexcitability and reduced seizure threshold.

Interestingly, while glutamatergic neuron morphology was unaffected by *dYEATS2* knockdown, behavioural assays revealed reduced locomotor function when *dYEATS2* was depleted specifically under the glutamatergic OK371-GAL4 driver (Figure 4I). The magnitude of impairment was milder than that observed with DAergic-specific knockdown and most prominent in young flies, suggesting that glutamatergic neurons may be functionally sensitive to dYEATS2 deficiency without overt structural degeneration.

These findings reinforce the notion that not all neuronal populations respond equally to transcriptional perturbations of dYEATS2; rather, vulnerability is shaped by intrinsic physiological demands and calcium-buffering capacity, manifesting as distinct degrees of cytotoxicity across cell types.

Given that downregulation of metabolic genes was also observed in the head transcriptome of *dYEATS2* knockdown flies (Figure S5), impaired energy metabolism likely represents an additional source of neuronal dysfunction. For instance, glutamatergic neurons, which appear morphologically resilient to calcium overload, could still be vulnerable to metabolic insufficiency due to their high energetic demands for sustained synaptic activity (Quintela-Lopez et al. 2022). Thus, while dopaminergic neurons could primarily succumb to calcium-driven excitotoxic stress, glutamatergic populations may experience dysfunction through a distinct, metabolism-based mechanism. These complementary vulnerabilities underscore the multifaceted consequences of dYEATS2 deficiency and illustrate how epigenetic perturbation can differentially compromise neuronal subtypes depending on their physiological and metabolic requirements.

### Targeting SOCE confirms calcium-dependent vulnerability of DAergic neurons in *dYEATS2*-deficient flies

Given that *dYEATS2* depletion leads to elevated intracellular calcium and correlates with DAergic neuron loss, we investigated whether calcium overload directly drives this selective neurodegeneration. To test this, we pharmacologically attenuated SOCE using the Orai inhibitor BTP2, previously shown to improve neurobehavioral deficits in *dYEATS2*-deficient flies. We then assessed DAergic neuron structural integrity in adult flies expressing *mCD8-GFP* and *dYEATS2 RNAi* specifically in DA neurons (*Ddc>mCD8-GFP>dYEATS2 IR*; Figure 5A).

**Figure 5.**
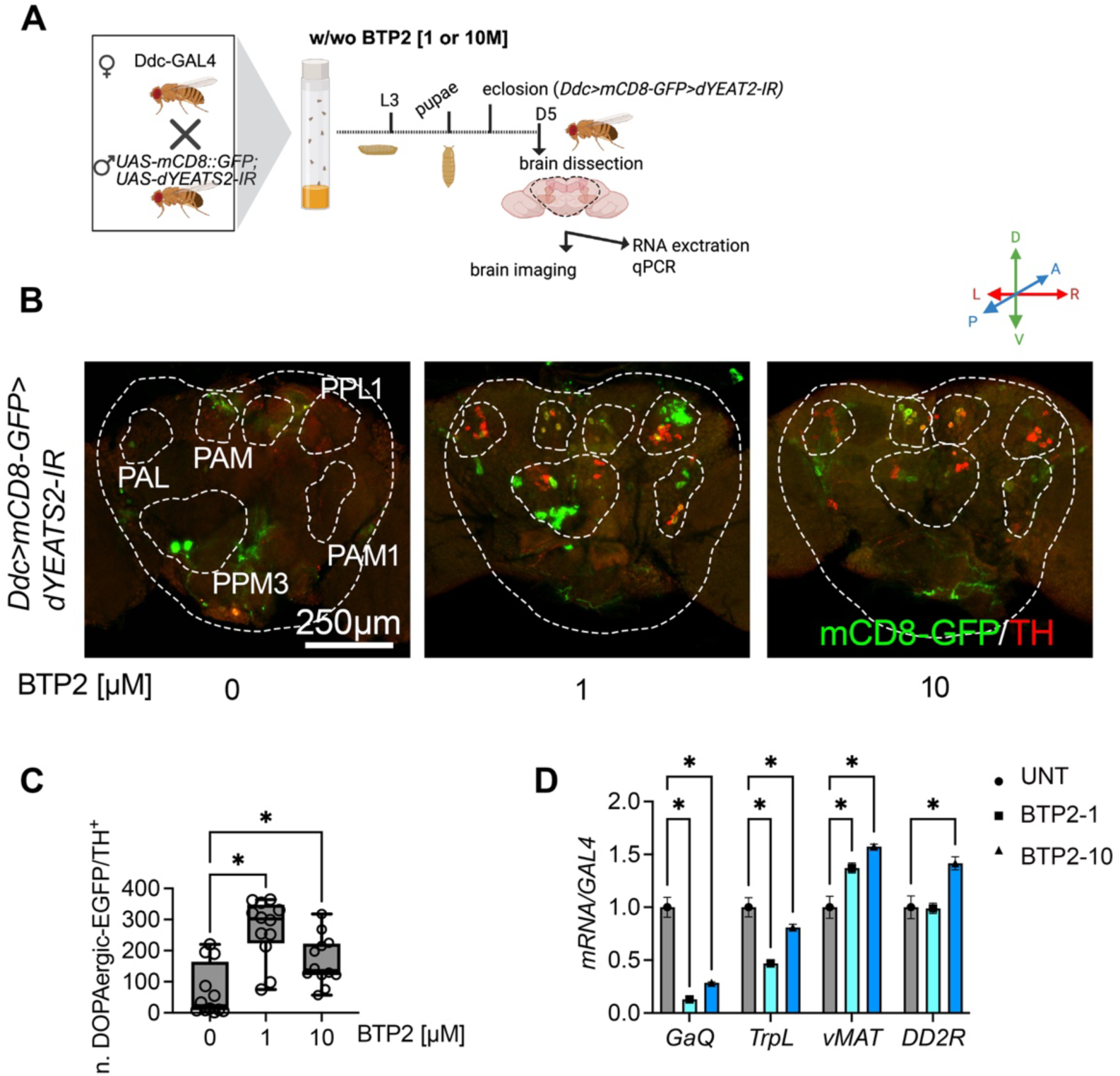
Pharmacological attenuation of SOCE restores DAergic synaptic integrity in *dYEATS2*-deficient flies. (A) Experimental scheme. Flies expressing membrane-tethered mCD8-GFP and *dYEATS2* RNAi specifically in dopaminergic neurons (*Ddc>mCD8-GFP>dYEATS2-IR*) were reared on standard medium supplemented with vehicle or the Orai inhibitor BTP2 (YM-58483) at 1 μM or 10 μM. Adult flies were transferred to fresh vials containing the same treatments and heads were dissected at 5 days post-eclosion for confocal imaging or RNA extraction. (B) Representative confocal images (posterior→anterior orientation) showing functionally active DAergic neurons identified by co-localization of mCD8-GFP (membrane marker expressed under Ddc-GAL4 driver) and Tyrosine Hydroxylase (TH) immunoreactivity. Central brain boundaries are indicated by dotted lines; major DA clusters (PAL, PAM, PPL1, PPM3) are highlighted with red dashed circles. (C) Quantification of EGFP– TH co-localization (number of co-localized puncta) was performed on 10 independent brains per condition using the colocalization module in CellSense (Olympus). Bars show mean ± SEM; BTP2 treatment at both 1 μM and 10 μM significantly increased the number of EGFP– TH co-localizing spots relative to untreated *dYEATS2*-IR animals. (D) Transcript levels of selected *dYEATS2*-responsive genes (*Gαq*, *trpL*, *vMAT*, *DD2R*) were measured from dissected adult heads following vehicle or BTP2 treatment to assess whether SOCE inhibition modulates these transcriptional changes. Gene expression was determined by reverse transcription quantitative PCR (RT-qPCR). Data are presented as mean ± SEM. Statistical significance was assessed by one-way ANOVA with Šidák’s and Tukey’s post hoc tests, respectively; *p* < 0.05 was considered significant.

Confocal analysis revealed that BTP2 treatment at 1 μM and 10 μM significantly increased the number of TH-positive DA neurons, measured by co-localization of mCD8-GFP and TH immunoreactivity (Figures 5B–C, S6). These results indicate that inhibiting Orai-mediated calcium influx mitigates the structural deficits caused by *dYEATS2* knockdown, establishing maladaptive calcium signalling as a dYEATS2-dependent causal factor in DAergic neuron degeneration.

BTP2 treatment not only restored DAergic neuron numbers but also normalized the expression of calcium signalling genes such as *GαQ* and *trpL*, which are upregulated in dYEATS2-deficient neurons (Figure 5D). These findings indicate a calcium-dependent feedback loop whereby elevated intracellular calcium drives sustained expression of calcium entry machinery and GPCR effectors. Inhibiting Orai with BTP2 disrupts this loop, reducing calcium influx and rebalancing transcription. Thus, dYEATS2 deficiency promotes maladaptive calcium signalling that reprograms gene expression and compromises neuronal viability, while Orai inhibition rescues both structural and transcriptional phenotypes.

Furthermore, BTP2 treatment normalized the expression of *vMAT* and *DD2R*, which were significantly downregulated in *dYEATS2*-deficient flies prior to treatment as shown in (Lo Piccolo et al. 2024) (Figure 5D). The recovery of these crucial DAergic pathway genes likely reflects restored neuronal function and integrity, as their expression depends on healthy DAergic neuron status.

Taken together, these findings indicate that calcium dysregulation is a central mechanism linking epigenetic disruption caused by dYEATS2 deficiency to selective DAergic neuron degeneration. They also show that targeted SOCE inhibition effectively restores calcium balance in this genetic context.

Beyond the acute calcium dysregulation elicited by dYEATS2 depletion, our results suggest that perturbations in crotonate metabolism may also influence calcium signalling in dopaminergic neurons. Because YEATS2 acts as a selective reader of histone crotonylation, its loss may mimic the functional consequences of diminished crotonylation by disrupting recognition of this chromatin modification. Accordingly, alterations in crotonyl-CoA abundance or in the enzymatic machinery that regulates histone crotonylation could similarly compromise calcium homeostasis, even in the absence of direct genetic mutations. Together, these observations suggest a novel mechanism whereby metabolic–epigenetic crosstalk converges on calcium signalling to shape dopaminergic neuron vulnerability, offering a conceptual framework for future studies aimed at dissecting selective neuronal susceptibility and identifying potential therapeutic avenues in neurodegenerative disease.

## Materials and Methods

### Drosophila husbandry

*Drosophila melanogaster* stocks were maintained on standard cornmeal–yeast–glucose medium at 22 °C with 60% humidity under a 12:12 light/dark cycle, unless otherwise specified. The GAL4/UAS binary system (Brands et al. 2024) was employed for tissue-specific knockdown or overexpression. Pan-neuronal manipulations were driven by nSyb-GAL4, while dopaminergic, glutamatergic, or mushroom body–specific expression was achieved using Ddc-GAL4, OK371-GAL4, and OK107-GAL4 drivers, respectively, at 25 °C. The full list of strains used in this study is provided in Table S2. To minimize genetic background effects, all experimental flies were backcrossed six times with the *w* strain before use.

### Library preparation and RNA sequencing (RNA-seq)

For transcriptomic analysis, 3-day-old male flies of each genotype were collected, snap-frozen in liquid nitrogen, and decapitated. Total RNA was extracted from 50 heads per replicate (three biological replicates per group) using the RNeasy Mini Kit (Qiagen, Valencia, CA) following the manufacturer’s instructions. The RNA samples, stabilized under ethanol precipitation conditions, were submitted to Macrogen, Inc. (Korea) for RNA sequencing (RNA-Seq). Prior to sequencing, the samples were analyzed with an Agilent 2100 Bioanalyzer to assess overall quality. Only samples with an RNA Integrity Number (RIN) > 7 and an rRNA ratio > 1 were processed further. RNA-sequencing libraries were prepared using the TruSeq Stranded mRNA Library Prep Kit, and sequencing was performed on the Illumina NovaSeq 6000 platform with a read length of 100 bp paired-end, generating approximately 30 million reads per sample.

### Quality control and preprocessing of RNA-seq data

Illumina short-read sequencing data were retrieved from Macrogen’s online repository and then transferred to the local server – MedCMU HPC – for analysis. Raw FASTQ files were processed using the nf-core/rnaseq pipeline v3.14.0 (Ewels et al. 2020) executed via Nextflow environments (Di Tommaso et al. 2017). Specifically, per-sample quality control was performed with FastQC v0.12.0; adapter trimming was conducted with Trim Galore! v0.6.5 (Krueger 2015); putative genome contaminants were removed using BBSplit (Bushnell 2014); and rRNA was depleted *in silico* with SortMeRNA (Kopylova et al. 2012). Clean reads were aligned to the *Drosophila melanogaster* reference genome BDGP6 (dm6) (Hoskins et al. 2015) using HiSat2 (Kim et al. 2019). The resulting BAM files were coordinate-sorted and indexed using SAMtools (Li et al. 2009). Duplicate molecules in UMI-tagged libraries were collapsed with UMI-tools (Smith et al. 2017), while duplicates in non-UMI libraries were marked with Picard MarkDuplicates. Summary metrics of all replicates from different tools were compiled with MultiQC V1.19 (Table S1). Reference-guided quantification was performed with StringTie2 (Kovaka et al. 2019) against the Ensembl release 81. Count data generated by StringTie2 in quantification-only mode were then used as input for differential expression analysis with the DESeq2 package (Love et al. 2014) in the R software environment v4.1.

### Functional enrichment analysis

The matrix of DESeq2 normalized read counts (median of ratios) was used as an input of expression dataset for Gene Set Enrichment Analysis (GSEA) analysis, which was performed with GSEA software version 4.3.2 on predefined gene sets derived from 137 KEGG pathways in *D. melanogaster* (Cheng et al. 2021). To identify concordantly regulated pathways between the knockdown and control groups, GSEA was conducted using the following parameters: permute = gene set, metric = Signal2Noise, and scoring_scheme = weighted (Song et al. 2023), with the Kolmogorov-Smirnov statistic. Gene sets with a false discovery rate (FDR) < 0.05 were considered significantly deregulated and enriched gene sets. Leading-edge analysis was then performed on the considerably enriched gene sets in GSEA. A Heat map of candidate gene lists and a bubble plot based on the GSEA analysis were plotted by an online platform for data analysis and visualization (http://www.bioinformatics.com.cn/srplot).

### RNA extraction and RT-qPCR

Total RNA was extracted from Drosophila tissues using the RNeasy® Mini Kit (QIAGEN) according to the manufacturer’s instructions. Three independent biological replicates were prepared. RNA quality and concentration were measured with a NanoDrop One/OneC spectrophotometer (Thermo Fisher Scientific). For cDNA synthesis, 1 µg of RNA was reverse transcribed using the SensiFAST cDNA Synthesis Kit (Bioline). Quantitative RT-PCR (RT-qPCR) was performed in technical triplicates for each biological replicate with the SensiFAST SYBR® Lo-ROX Kit on a CFX Opus Real-Time PCR System (Bio-Rad), employing gene-specific primers (Table S3). ACTIN was used as a reference gene for normalization.

### Protein extraction and western blotting

Protein extracts were obtained from adult *Drosophila* heads using RIPA Lysis and Extraction Buffer (Thermo Fisher) supplemented with protease inhibitors. Approximately 40 heads were homogenized in 100 µl of buffer, followed by incubation at 4 °C with gentle agitation for 20 minutes. Lysates were cleared by centrifugation at 14,000 × g for 20 minutes at 4 °C, and protein concentrations were determined using the BCA Protein Assay Kit (Millipore #71285). Equal amounts of protein (30 µg) were separated on 12% polyacrylamide gels and transferred to Immobilon-P PVDF membranes (Merck) with the Trans-Blot Turbo transfer system (Bio-Rad). Membranes were blocked in 5% skim milk prepared in TBS-T (Tris-buffered saline, 0.1% Tween-20) and incubated overnight at 4 °C with the indicated primary antibodies (Table S4). After four washes with TBS-T, membranes were exposed to HRP-conjugated secondary antibodies (1:3000) for 1 hour at room temperature. Both primary and secondary antibodies were diluted in 5% skim milk/TBS-T. Signals were visualized using WesternSure chemiluminescent reagents (LI-COR) and captured with a C-DiGit Blot Scanner (LI-COR). Protein expression was normalized to Actin, and band intensities were quantified with ImageJ (v1.53k) across independent biological replicates. Additional information on chemicals is listed in Table S5.

### Brain dissection and imaging

Brains from 5-day-old adult flies carrying the pan-neuronal driver and UAS-GCaMP6s were dissected in ice-cold 1× PBS. Following dissection, tissues were incubated *ex vivo* for 20 min at 25 °C in PBS supplemented with either 200 mM NaCl, 200 mM KCl, or 200 mM glucose. After incubation, brains were mounted on glass slides using antifade mounting medium and imaged for GFP fluorescence. For analysis of specific neuronal populations and mushroom body architecture, brains from 5-day-old adults were dissected in Schneider’s *Drosophila* medium and fixed in 4% formaldehyde prepared in PBS for 30 min. Samples were then rinsed three times for 20 min each in PBS containing 0.3% Triton X-100, followed by blocking in PBS with 0.3% Triton X-100 and 0.2% bovine serum albumin for 30 min. Primary antibodies, mouse anti-FasII (1D4, Developmental Studies Hybridoma Bank) or rabbit anti-TH (AB152, Merck), were applied in blocking buffer and incubated for 48h at 4 °C. Brains were subsequently washed and incubated overnight at 4°C with Alexa Fluor-conjugated secondary antibodies (see Table S4). Images were acquired using an Olympus Fluoview FV3000 confocal microscope equipped with a Plan Apo 20×/0.80 objective lens. Identical laser and detector settings were used across experimental groups to ensure comparability. Z-stacks were collected at 2048 × 2048-pixel resolution, with 12 optical sections (4 µm step size), and processed as maximum-intensity projections oriented from posterior to anterior.

### Quantification of Dopaminergic Neuronal Clusters and Colocalization Analysis

To measure the extent of GFP-labeled neuronal populations, flies expressing UAS-mCD8-GFP under the control of specific neuronal drivers were analyzed. Maximum projection images were generated from confocal stacks using CellSense Dimension software (version 3.2, Olympus). Threshold levels for object detection were established using control samples (*mCD8-GFP; NTC-IR*), and the same parameters were subsequently applied to experimental genotypes (*mCD8-GFP; dYEATS2-IR*). Quantification was performed within defined regions of interest (ROIs), and automated object detection and measurement were used to calculate the total number of GFP-positive neurons per ROI. To assess the subset of DAergic neurons with functional activity, we quantified colocalization between GFP and TH immunoreactivity. In this context, GFP marked neurons expressing UAS-mCD8-GFP under control of the Ddc-GAL4 driver, while TH served as a definitive marker of dopamine-producing neurons. Confocal z-stacks were imported into CellSense Dimension software (version 3.2, Olympus), and colocalization analyses of GFP and TH channels were performed across 12 optical sections (4 µm each). ROIs were drawn around the central brain lobes. Scatterplots of the two channels were generated, and colocalization files were created for each ROI. Segmentation was applied with a minimum object size of 50 pixels, followed by automatic thresholding to standardize detection. Quantification of GFP-positive, TH-positive, and colocalized (GFP⁺/TH⁺) neurons was then performed to determine the proportion of functionally dopaminergic cells.

### Seizure-like behaviours

Mechanical stress–induced seizure susceptibility was assessed using the bang-sensitivity assay, adapted from previously described protocols (Lo Piccolo et al. 2024). Newly eclosed males were aged for 3–4 days and gently transferred individually into empty vials using a suction pipette to avoid CO₂ anesthesia. After a 5-min acclimation period, flies were subjected to a 10-s high-speed vortex. Immediately thereafter, they were placed in a 15-cm glass Petri dish and video-recorded for offline analysis. Seizure-like behavior (SLB) was defined as uncontrolled movements or paralysis persisting for more than 3 s after mechanical stimulation. Recovery time was recorded, and values were compared with genotype-matched controls. For each genotype, a minimum of 50 flies was analysed.

### Climbing assay

Adult locomotor performance was evaluated using a climbing paradigm adapted from (Jantrapirom et al. 2018). Male flies of each genotype were collected at eclosion and aged for 3, 9, 15, 21, or 27 days. Groups of ten age-matched flies were transferred, without anaesthesia, into clear conical tubes and allowed to acclimate for 10 min. To initiate climbing, tubes were gently tapped to displace the flies to the bottom; this stimulus was applied at 30-s intervals and repeated five times for each group. Climbing behaviour was video recorded, and performance was quantified by scoring the vertical distance reached within 5 s: 0 (<2.0 cm), 1 (2.0–3.9 cm), 2 (4.0–5.9 cm), 3 (6.0–7.9 cm), 4 (8.0–9.9 cm), and 5 (≥10.0 cm). A climbing index was calculated for each group by multiplying the number of flies in each category by its score, summing across categories, and dividing by the total number of flies. For each genotype, assays were performed on at least 100 flies, and values were compared with genotype-matched controls.

### Data analysis

All statistical analyses were performed using GraphPad Prism 9 (GraphPad Software). For datasets following a normal distribution, two-group comparisons were assessed with either Student’s *t*-test or Welch’s *t*-test, while differences among more than two groups were evaluated using one-way ANOVA with Tukey’s or Dunnett’s post hoc test. When data did not meet normality assumptions, the Mann–Whitney U test was applied for pairwise comparisons, and the Kruskal–Walli’s test with Dunn’s post hoc analysis was used for multiple groups. A *p*-value < 0.05 was considered statistically significant. Results are shown as mean ± standard deviation (SD) for experiments with fewer than six replicates (e.g., Western blotting and qPCR), and as mean ± standard error of the mean (SEM) for larger datasets such as behavioural assays.

## Acknowledgments

This work was funded by the Mid-Career Research Grant (N42A670768) from the National Research Council of Thailand, with additional support from the Fundamental Fund of Chiang Mai University (grant FF68/207569) and the Faculty of Medicine, Chiang Mai University (grant FACMED 139/2567). Partial funding was also provided by Genomics Thailand through the Health Systems Research Institute (HSRI; grant 68-060). We are grateful to Mr. Papon Hitmool, Mr. Sakorn Phongchankhiao, and Mr. Tawan Munwanna (Department of Pharmacology, Faculty of Medicine, Chiang Mai University), as well as Ms. Chansunee Panto (CMUTEAM, Faculty of Medicine, Chiang Mai University), for their invaluable technical and administrative assistance. We also thank the Bloomington Drosophila Stock Center for providing fly strains, and Science and Educational Company Limited (SCIED) for confocal scanning electron microscopy training.

## Author Contributions

Conceptualization: L.LP. and S.J.; Data curation: L.LP and S.J.; Formal Analysis: L.LP, R.Y., P.N., and S.J.; Funding acquisition; L.LP; Investigation: : L.LP, R.Y., P.N., and S.J.; Methodology: L.LP., R.Y. and P.N.; Project administration: L.LP; Resources; L.LP and S.J; Software: L.LP. and R.Y.; Supervision; L.LP; Validation: L.LP. and S.J.; Visualization: L.LP., R.Y. and S.J.; Writing – original draft: L.LP.; Writing – review & editing: All authors have reviewed and approved the final version of this manuscript.

